# Interspecies interaction reshapes the fitness landscape of evolved genotypes

**DOI:** 10.1101/2024.07.06.602328

**Authors:** Xinli Sun, Zhihui Xu, Guohai Hu, Jiyu Xie, Yun Li, Lili Tao, Nan Zhang, Weibing Xun, Youzhi Miao, Ruifu Zhang, Qirong Shen, Ákos T. Kovács

**Author notes:** Xinli Sun and Zhihui Xu contributed equally to this work. Corresponding authors: Zhihui Xu, Qirong Shen, and Ákos T. Kovács.

## Abstract

Microbial interaction and their evolution is vital for shaping the structure and function of microbial communities. However, the mechanisms governing the directionality and stability of the evolution of interactions within microbial communities remain poorly understood. Here, we used a syntrophic two-species biofilm consortium composed of *Bacillus velezensis* SQR9 and *Pseudomonas stutzeri* XL272 that promotes plant growth through their metabolic interactions and investigated how the interactions within the consortium change over evolutionary timescale to characterize the phenotypic and genetic diversification. The focal species *B. velezensis* SQR9 rapidly diversified into diverse colony morphotypes, both in the presence and absence of its interactor, *P. stutzeri* XL272, with variable frequencies. These morphotypes displayed phenotypic differentiation among biofilm formation, planktonic growth, and spore formation. The evolved *P. stutzeri* altered the fitness landscape of *B. velezensis* morphotypes, allowing the weaker rough morphotype to outcompete the biofilm-enhanced slimy morphotype. Whole genome re-sequencing correlated these phenotypic changes with mutations in specific genes encoding regulators of *B. velezensis*, including *ywcC*, *comA*, *comP*, *degS*, *degQ* and *spo0F*. The coevolutionary partner, *P. stutzeri* increased its exopolysaccharide production that could be explained by a frame shift mutation in *cpsA* gene encoding capsular polysaccharide (CPS) biosynthesis protein. Compared with the mono-evolution, co-evolved *B. velezensis* populations showed greater mutation accumulation in intergenic regions, which led to greater genetic parallelism. Furthermore, the dissimilarity between mono-evolved and co-evolved populations increased over time. Our study reveals intricate genetic diversification and fitness differentiation within a biofilm consortium, shaped by both abiotic conditions and biotic interactions.

## Introduction

Evolutionary diversification rapidly emerges even when bacteria adapt to a new environment in isolation ^1–3^. Intraspecific competition, heterogenous environment ^2,3^, eco-evolutionary feedbacks ^4^, trade-offs ^5^ and clonal interference ^6^ are key drivers of adaptive diversification. The intricate web of interactions between microorganisms, ranging from symbiotic to antagonistic interactions, is critical to shaping the composition and function of microbial communities. Several studies have documented that microbial interactions alter evolutionary response to new environments ^7–12^. Coevolution, the reciprocal evolutionary change between interacting species, is likely to shift the nature of interaction and change community-level properties ^13,14^. Interspecies interactions could promote the evolution of cross-feeding ^15,16^, adaptive function loss ^8,11,17^, niche expansion ^9^, but could also slow down evolution ^18,19^. In this context, elucidating the interplay between microbial interactions and evolutionary strategies represents a major challenge for modern biology. Despite the importance of eco-evolutionary dynamics and their impact on community properties, our knowledge is still limited on how evolution shift the balance of mutualism and competition in structured environments.

Investigating microbial evolution within two-species communities allows a focused dissection of the molecular mechanisms governing ecological feedback and evolutionary trajectories, which may be obscured in more diverse communities. In the present study, mono-evolution of *Bacillus velezensis* SQR9 and co-evolution between *B. velezensis* SQR9 and *Pseudomonas stutzeri* XL272 were investigated. In our previous research, we have demonstrated that these two soil-derived isolates form synergistic dual-species pellicle biofilm at 24h and benefit plant growth when co-inoculated to the soil ^20^. However, if the biofilm was collected at 48h, their interaction changed from mutualism to ammensalism. Comparing the phenotypic and genetic outcomes of single-species and dual-species pellicle evolution, we aimed to unravel how eco-evolutionary feedback changes interspecies interaction.

Under certain selective pressure, the evolutionary outcome is highly repeatable across species and environments, suggesting the possibility to forecast the evolution of related species in similar environments ^21^. Based on the previous biofilm experimental evolution reports from *Pseudomonas* spp. ^22–24^, *B. subtilis* ^25^, *B. thuringiensis* ^26^ (Lin et al 2022), and co-evolution of *Paenibacillus amylolyticus and Xanthomonas retroflexus* ^10^, we expected the following evolutionary outcomes from *B. velezensis – P. stutzeri* biofilm co-evolution: (1) Phenotypic trade-off, such as biofilm formation, public good production, motility, and growing capacity; (2) Mutations arise related to regulating cell adhesion, biofilm formation, sporulation, nutrient utilization, and so on; (3) *P. stutzeri* affect the evolution of *B. velezensis*, either by changing the evolutionary strategies, or by changing the fitness of different newly evolving genotypes. These hypotheses were tested by comparing the phenotypic and genotypic differences between ancestors and evolved genotypes, both in monoculture and coculture conditions.

## Results

### Evolved biofilm community increased biomass and exhibited diversification

To investigate the effect of interspecies interaction on bacterial evolution, we performed mono experimental evolution of *B. velezensis* SQR9 and coevolution of *B. velezensis* SQR9 – *P. stutzeri* XL272 consortium in pellicles (see Methods). The populations were diluted and transferred to fresh medium every other day (Fig 1A). Each transfer corresponded to approximately 6 to 7.42 generations. In our previous study, the pellicles were collected at 24 hours ^20^, however, the pellicles were collected at 48 hours to provide adequate time for biofilm co-culture maturation before serial transfer. From 24 to 48 hours, the interaction between *B. velezensis* and *P. stutzeri* changed from mutually beneficial cooperation to ammensalism (Fig S1).

**Figure 1.**
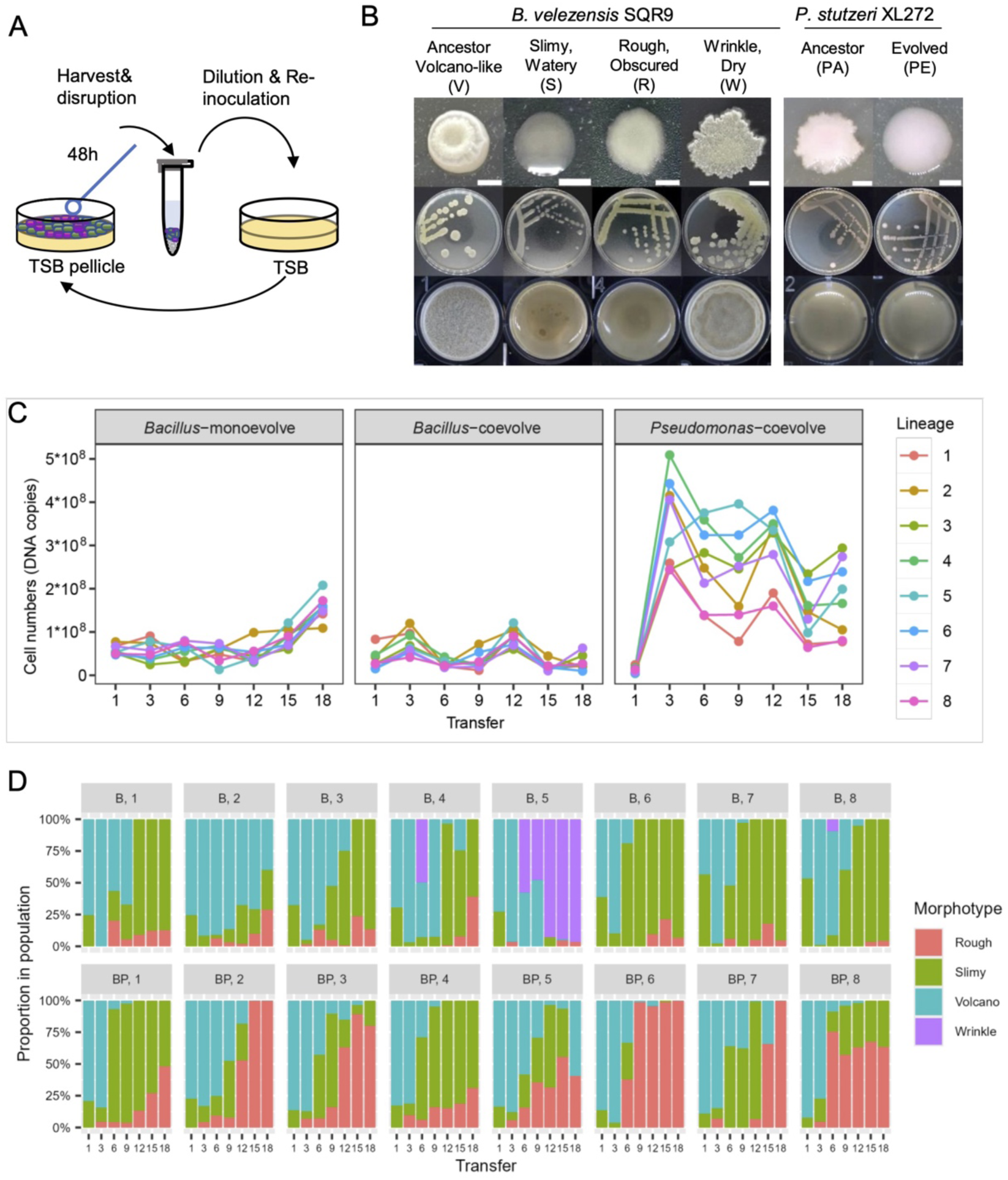
Population dynamics. **(A)** Evolutionary dynamics of mono-evolved *B. velezensis* populations. **(B)** Colony phenotype of evolved variants. Colonies with different morphotypes were streaked on TSB agar plates and cultivated in 30 °C. Petri dish diameter is 9 cm. Scale bar = 0.2 cm. **(C)** Evolutionary dynamics of co-evolved communities. Population or community productivity was assessed every 3rd transfer by qPCR. **(D)** The emergence and successional dynamics of *B. velezensis* evolved variants. “B” represents mono-evolved populations, “BP” represents co-evolved communities.

Pellicle appeared dry, frosted, and fragile at the onset of evolution (Fig S2). Pellicles became watery and glutinous at 6^th^ transfer and kept such a feature until the endpoint of the experimental evolution setup. Co-evolved pellicles were thicker and stickier than mono-evolved pellicles in general. The dynamic changes of cell numbers in pellicle populations were assessed by qPCR. Cell numbers of all *B. velezensis* mono-species biofilm populations increased gradually starting from 12^th^ transfer (Fig 1C). Population B5 yielded the highest biomass, while the biomass of population B2 was the lowest at the final transfer. Mono-evolution of *P. stutzeri* XL272 was not performed due to the thin and fragile pellicle formation of *Pseudomonas* cultures prohibiting harvesting the biofilm for the subsequential cultivation. In co-evolved biofilm communities, the cell numbers of *B. velezensis* maintained steadily (Fig 1C). The cell numbers of co-evolved *P. stutzeri* populations rapidly increased at the 3^rd^ transfer, then decreased to a stabilized level. The overall cell numbers of the co-evolved communities were higher than the mono-evolved populations.

The colony morphology was examined every 3^rd^ transfer on selective agar plates to identify phenotypic variants (Fig 1B) and their successional dynamics (Fig 1D). The ancestral *B. velezensis* SQR9 exhibits white, architecturally complex, three-dimensionally growing colony morphology, with a dry bulge and slimy edge, denoted as “Volcano” type. Three colony morphotype variants were identified: "Slimy" displayed a watery and glutinous colony, "Rough" exhibited a rough and obscured colony, while "Wrinkle" exhibited an enhanced wrinkled structure. The wrinkle morphotype only emerged in mono-evolved populations and only persisted in one population (B5). The ancestor-like volcano morphotype was gradually replaced by the variants and only persisted in few of the final populations (BP5, B2). In *P. stutzeri* XL272, the ancestral colony is irregular, while the evolved colonies are round. However, this difference is not discernible if the colonies are small, thus all the evolved clones are referred to as “PE”.

Interestingly, although the morphotype differentiation of *B. velezensis* was similar in mono-evolution and co-evolution, the successional dynamics were different – the proportion of rough morphotype within the evolving lineages was higher during co-evolution than during mono-evolution (Fig 1D). During the final transfer, 6/8 mono-evolved populations contained a large proportion of slimy types and a small proportion of rough types. In co-evolved populations, however, the proportion of rough greatly increased in 4/8 populations, and took over 3/8 populations entirely. A possible explanation for the increased occupation of rough type is the influence of *P. stutzeri*.

### Evolved clones displayed phenotypic differentiation

Next, we compared the phenotypic differences of the evolved clones, including growth curve, spore formation, and biofilm formation. Eight *B. velezensis* isolates were selected, either from mono-evolution or coevolution, with either slimy or rough morphotype. Comparing the growth kinetics under shaken condition showed that seven of the selected isolates increased growing capacity in shaken culture compared to ancestor, while six of the selected isolates displayed reduced growth rate (Fig S3A&B). Only the ancestor had the ability to form spores in pellicles at 2d and 7d, while no spores were observed in the evolved isolates (Fig 2B). Heat treatment followed by serial dilution and colony counting were used to determine the sporulation rate. While the sporulation rate of the ancestor was 2.0±0.27% and 85±6.4% for 2d and 7d, respectively, the evolved isolates displayed a spore frequency below 0.1%. In monoculture, compared with the ancestor, the slimy morphotype displayed an increased cell numbers in pellicles, while the rough morphotype had lower cell number in the biofilm (Fig 2C). No discernible changes were observed between *P. stutzeri* ancestor and the evolved variant regarding pellicle formation (Fig 1B & 2C).

**Figure 2.**
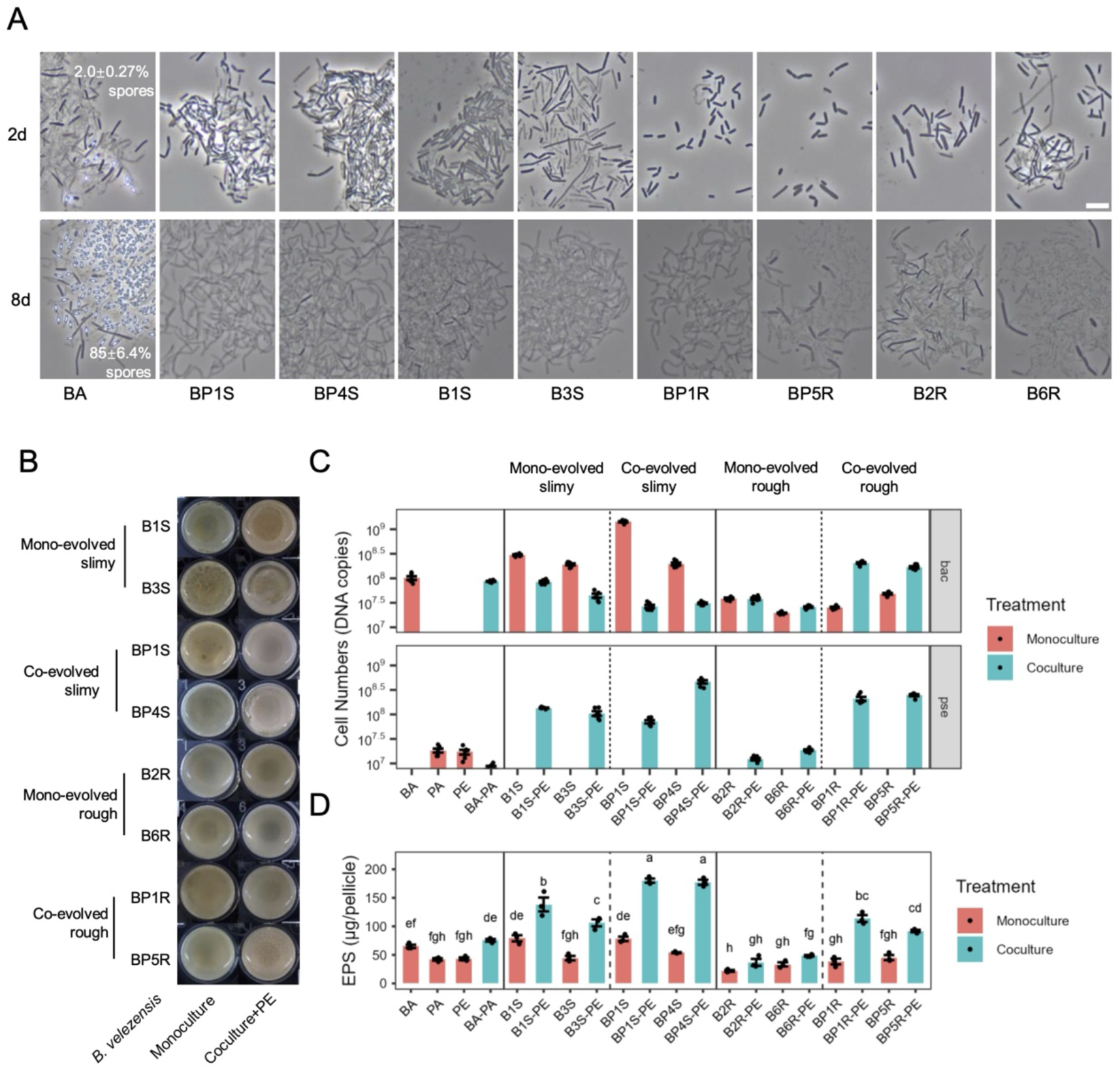
Comparison of ancestors and evolved isolates. **(A)** Representative images of sporulation in pellicles. Samples were collected 2 days and 7 days post incubation, and votexed rigorously. Scale bar represents 5 μm. No spores were observed except ancestor. **(B)** Pellicle morphotype. **(C)** Cell numbers quantification by qPCR. n = 6. The upper panel shows the cell numbers of *B. velezensis*, the down panel shows the cell numbers of *P. stutzeri*. **(C)** EPS quantification in pellicles. n = 3. Data shows the mean. The compact letter display indicates significant differences between groups based on Tukey’s HSD test (*p* < 0.05). “BA” represent *B. velezensis* ancestor, “PA” represents *P. stutzeri* ancestor. “Bn” represents monoevolved lineage, “BPn” represents coevolved lineage, “S” represents slimy morphotype, “R” represents rough morphotype.

To examine if evolutionary diversification affected bacterial interaction, we compared the dual-species pellicle formed by the ancestral *B. velezensis* and *P. stutzeri* strains with the evolved isolates of the two species. One *P. stutzeri* evolved isolate PE4 was selected as a representative, denoted as “PE”. Dual-species pellicles produced by co-evolved *B. velezensis* isolates were thicker than those produced by monocultures of the isolates (Fig 2B). We compared the total numbers of cells within the pellicles by qPCR. Interestingly, although the coevolved rough isolates of *B. velezensis* formed weaker pellicle than the coevolved slimy isolates in monoculture, cell numbers were increased in their coculture with *P. stutzeri*, while the slimy isolates were inhibited (Fig 2C). In most cases, *P. stutzeri* benefited from coculturing with *B. velezensis* evolved isolates (Fig 2C). Based on these results, we interpreted that the coevolved rough isolates of *B. velezensis* had a disadvantage during mono-evolution, while it had competitive advantage during co-evolution. Co-evolution was a prerequisite to enhance dual-species pellicle formation between the two species.

Since most pellicle lineages evolved to be slimy, we hypothesized that evolved isolates would produce more exopolysaccharide (EPS) than their ancestors. Comparing *B. velezensis* evolved isolates with their ancestors in mono-species biofilms, they produced similar or lower levels of EPS (Fig 2D). In general, dual-species biofilms produced higher levels of EPS compared to mono-species biofilms, except for those containing mono-evolved rough velezensis isolates. Based on the observation that the slimy isolates of *B. velezensis* decreased cell numbers, the evolved *P. stutzeri* isolates could be responsible for the increased EPS.

Taken together, these results suggested that the evolved *B. velezensis* morphotypes displayed phenotypic differentiation: although the evolved isolates increase growing capacity in shaken culture, they have a decreased growth rate and are incapable of sporulation. The morphotypes differ in their ability to form pellicles: slimy isolates of *B. velezensis* have a benefit in mono-evolution, whereas rough isolates have a benefit in coevolution. Co-evolution enhances biofilm formation by increasing both EPS production and the density of cells.

### Rough isolates exploit the matrix in *B. velezensis* populations, but cooperate in a dual-species consortium

We tested whether the pellicle formation of the coevolved rough isolates could be complemented by the extracellular matrix produced by the slimy *B. velezensis* isolates or the supernatants of the evolved *P. stutzeri* isolate. The rough isolates improved pellicle formation when supplemented with ancestor’s or slimy isolates’ extracellular matrix, but biofilm matrix from itself had no influence (Fig 3). This result suggests that rough isolates act as exploiters in *B. velezensis* mono-species populations. In addition, the supernatant of evolved *P. stutzeri* can also stimulate pellicle formation in rough isolates. As both rough isolates and evolved *P. stutzeri* benefited from coculture, their interaction is bidirectional cooperation. Therefore, the rough isolates had an increased fitness in the evolved biofilms by exploiting the extracellular substances released by its siblings or co-existing species.

**Figure 3.**
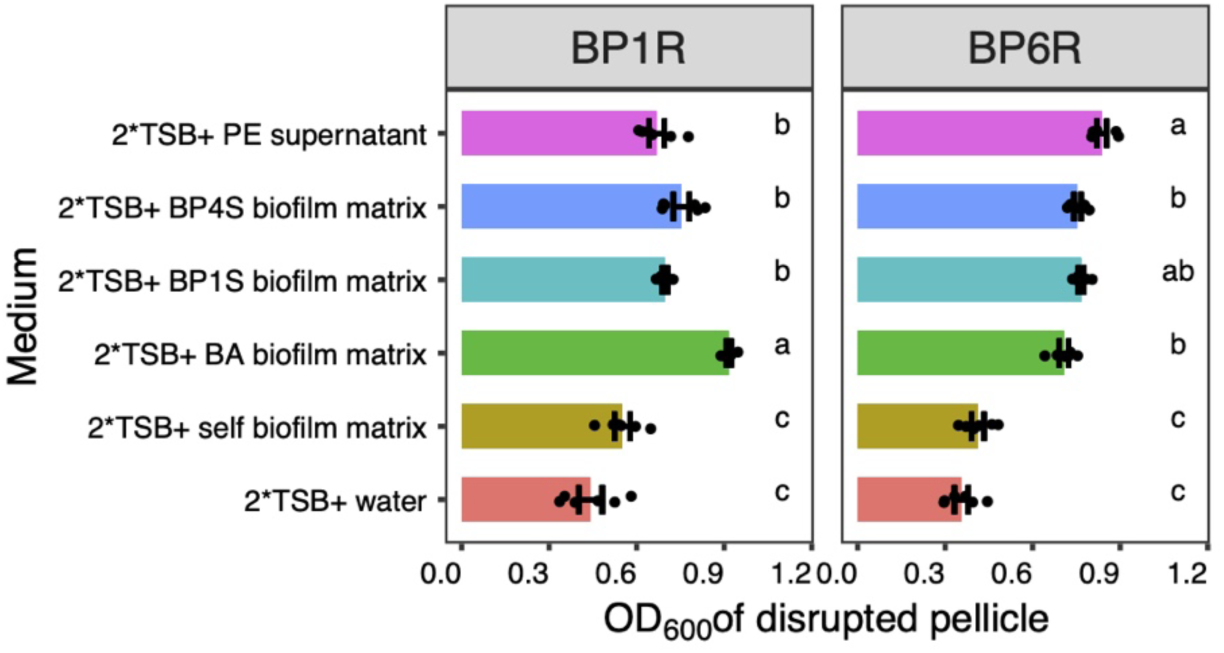
Pellicle complementation of rough isolates by extracellular matrix of other isolates. Different letters indicate significant difference (*p*<0.01) based on one-way ANOVA, tukey’s posthoc test. Pellicle biomass was measured by OD_600_ of votexed pellicle.

### Phenotypic differentiation correlates with certain mutations

To understand the genetic bases of phenotype differentiation and to distinguish mutations adapted to culture condition versus those responding to the presence of another species, the genomes of 69 *B. velezensis* evolved clones (16 volcano types, 25 slimy types, 26 rough types, 2 wrinkle types) and 16 *P. stutzeri* evolved clones were sequenced from the two evolutionary setups. Additionally, the genomes of the two ancestors were also re-sequenced to screen for mutations that emerged before the evolution experiment. 17 common mutations were detected in the ancestral *B. velezensis*, 7 mutations in the ancestral *P. stutzeri* compared with the genome sequence available in NCBI (Table S2), which were excluded from further analysis. Focusing on mutations present in at least two clones (Fig. 4 & S4), we observed that in *B. velezensis*, the most frequently mutated genes were *ywcC* that encodes a regulator controlling the expression of the *slrA* gene, and the *spo0F* gene that encodes phosphotransferase of the sporulation initiation phosphorelay. The *ywcC* mutation leads to depression of *slrA* transcription, thereby SlrR/SlrA stimulates biofilm formation by antagonizing the global negative regulator of biofilm development, SinR ^27^. These two genes harbored missense single nucleotide polymorphisms (SNPs) or frameshift mutations at different positions. We constructed knockout mutants of these two genes, the Δ*spo0F* displayed slimy colony phenotype with reduced pellicle formation, whereas the Δ*ywcC* mutant, though morphologically comparable to the ancestor, displayed enhanced pellicle formation (Fig 4B & C). These findings suggest that the slimy morphotype change is likely due to the *spo0F* mutation, while the enhanced pellicle formation can be attributed to the *ywcC* mutation.

**Figure 4.**
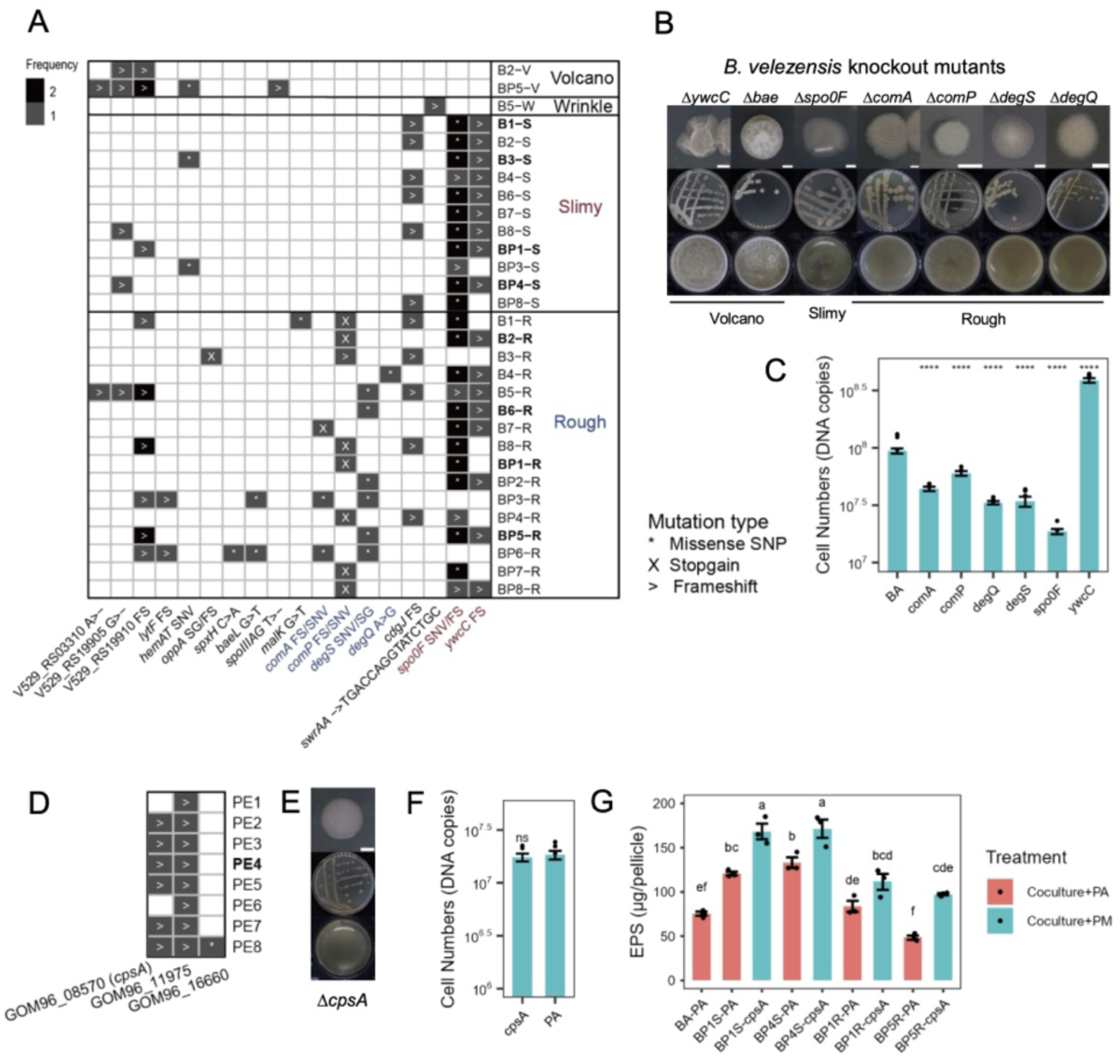
Common mutations identified in evolved clones from 18^th^ transfer and representative mutants. **(A)** *B. velezensis* SQR9 mutations. Isolates in bold text were selected for phenotypic characterization. V abbreviates for volcano, W abbreviates for wrinkle, S abbreviates for slimy, R abbreviates for rough. **(B)** Colony and pellicle morphotype of *B. velezensis* mutants. Scale bar represents 0.2 cm, Petri dish diameter is 9 cm, well diameter is 1.5 cm. **(C)** Cell numbers quantification of pellicles by qPCR. n = 6. “****” indicates significant difference compared to “BA” (t test, *p* < 0.01). **(D)** *P. stutzeri* XL272 mutations. PE abbreviates for evolved *P. stutzeri*. For both (A) & (D), “n” indicated the population. “*” and SNP refer to missence single nucleotide polymorphism, “>” and FS refers to frameshift mutation, “X” and SG refers to stopgain mutation. All the isolates shown above were isolated from the 18^th^ transfer. Isolates from the 9^th^ transfer were shown in Fig S4. **(E)** Colony and pellicle morphotype of *P. stutzeri ΔcpsA* mutant. **(F)** Cell numbers quantification of pellicles by qPCR. n = 6. “ns” indicates no significant difference compared to “PA” (t test, *p* < 0.01). **(G)** Exopolysaccharide (EPS) quantification of pellicles. n = 3. Different letters indicate significant differences based on ANOVA, tukey’s posthoc test (*p* < 0.05).

The rough morphotype carried mutations either in the *comA*/*comP* genes or *degS*/*degQ* genes. The ComP-ComA two-component signal transduction system regulates quorum sensing and competence development in *Bacillus subtilis* species ^28,29^. The sensor kinase DegS, together with its response regulator DegU, co-ordinate multicellular behavior including biofilm formation, genetic competence, motility, degradative enzyme production, and poly-γ-glutamic acid production ^30^. DegQ protein enhances the phosphate transfer from DegS to DegU, deletion of the *degQ* gene impairs complex colony architecture and biofilm formation ^31^. Knockout mutants of these four genes, *comA*/*P* and *degS*/*Q*, all displayed rough colony phenotype and reduced pellicle formation (Fig 4B & C). The two wrinkle clones shared frameshift mutation in the *swrAA* gene that encodes the swarming motility protein ^32^.

In *P. stutzeri*, only three parallel mutations were identified in at least two clones (Fig 4D). One of which was shared in all clones, a frameshift insertion in the GOM96_11975 gene encoding histidine kinase that regulates unknown function. This mutation was likely acquired in the overnight culture of the ancestor used to initiate the evolution experiment. Most clones shared an identical frameshift insertion at the high C region in the GOM96_08570 gene that encodes a CPS biosynthesis protein, we designated as *cpsA*. Given that the evolved co-culture pellicles became slimy, we hypothesized a correlation between this gene with polysaccharide production. The production of CPS was reported to interfere with biofilm formation across various bacteria ^33–35^. Thus, we constructed a complete knockout strain Δ*cpsA* (Fig 4E). The colony phenotype of Δ*cpsA* mimicked the evolved *P. stutzeri* clone with a round colony edge (Fig 2C). Pellicle formation did not significantly differ between PA and Δ*cpsA* (Fig 4F). Notably, the dual-species biofilm containing the Δ*cpsA* mutant showed higher EPS production compared to the ancestor (Fig 4G). This effect was not due to an increase in biomass, as measured by OD_600_, indicating that the disruption of the *cpsA* gene is responsible for the higher levels of EPS.

Taken together, our genomic analysis reveals that specific mutations drive phenotype differentiation. In *B. velezensis*, *spo0F* mutations were linked to a slimy morphotype change, while mutations in *comA*/*comP* and *degS*/*degQ* were associated with rough morphotypes. Additionally, *ywcC* mutations were linked to enhanced biofilm formation. In *P. stutzeri*, *cpsA* mutations increased exopolysaccharide production in dual-species biofilms.

### Differential population genetic dynamics of biofilm evolution

Using longitudinal population sequencing of all lineages at every 3^rd^ transfer (and also the 1^st^ transfer), we compared the mutational dynamics of mono-evolution to co-evolution. Co-evolved populations of *B. velezensis* contain more mutations than mono-evolved populations (Fig 5A), and most of these mutations are intergenic (Fig 5B). The Jaccard similarity coefficient (*J*) of all pair of lineages from each evolution group was calculated. The higher the value, the more similar the two lineages are. It is calculated by dividing the number of shared mutations by the number of total mutations of two independent lineages. Interestingly, the *J* values of co-evolved populations are significantly higher than those of mono-evolved populations (*p* < 0.001) (Fig 5C), indicating a greater parallelism for co-evolved populations. Excluding mutations in intergenic regions from analysis showed no significant difference in *J* values between mono-evolved and co-evolved lineages (Fig S5A). This suggests that the observed high parallelism in co-evolution could be attributed to parallel mutations specifically in intergenic regions.

**Figure 5.**
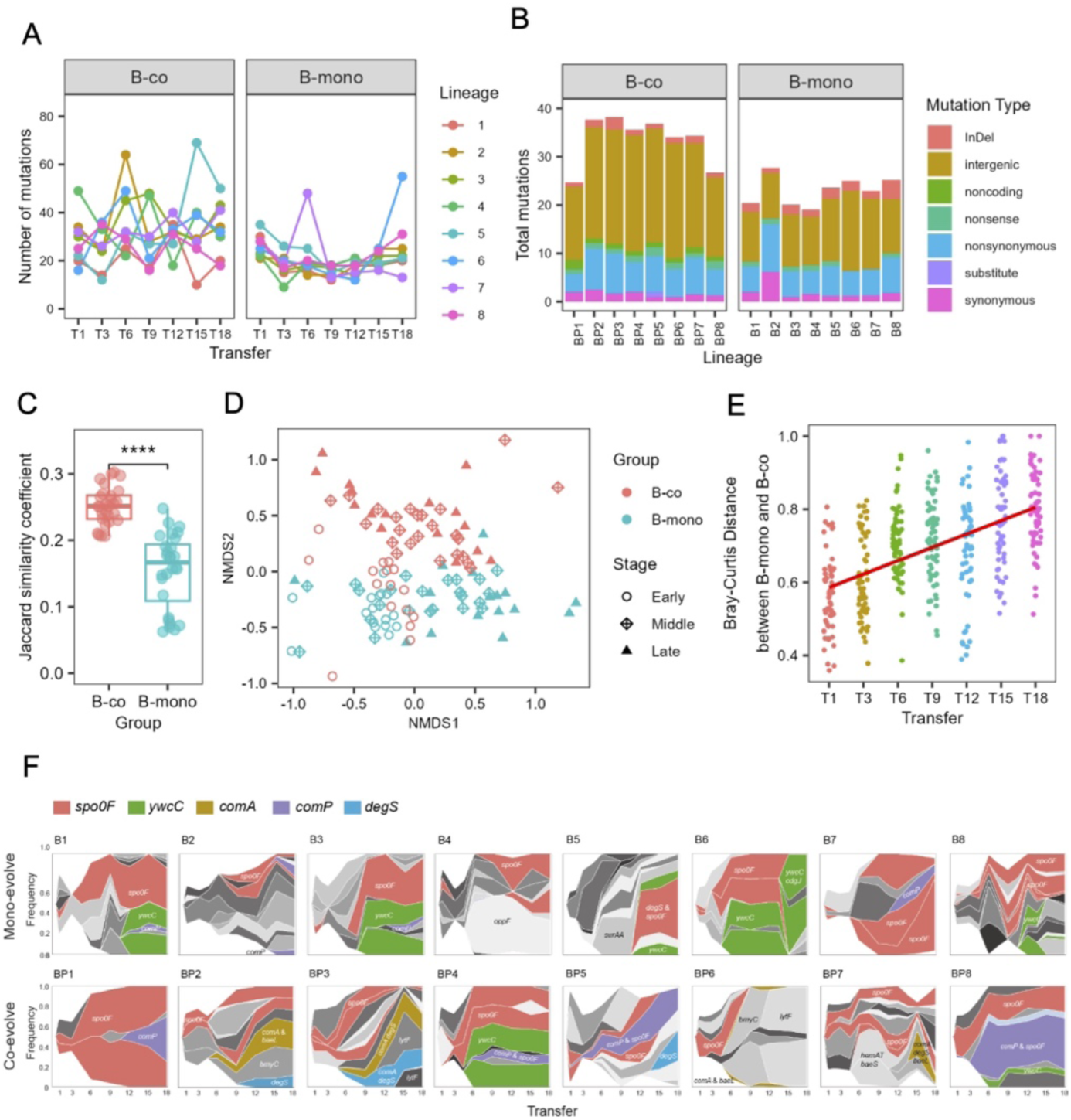
Evolutionary dynamics of populations. **(A)** Dynamic distribution of mutations detected in each lineage over time. **(B)** Distribution of mutation type of each lineage. **(C)** Degres of parallelism within each group estimated by Jaccard index. Asterisks indicate significant differences between co-evolved and mono-evolved *B. velezensis* populations (t test, \*\*\**p* < 0.001). Dots indicate the J value of paired-wise comparison between lineages. The Jaccard index describes the likelihood that the same gene is mutated in two independent lineages and ranges from 0 (no mutated genes in the two lineages are shared, G1∩G2 = 0) to 1 (the two lineages have exactly the same set of mutated genes, G1 = G2). **(D)** Non-metric multidimensional scaling (NMDS) ordinations of co-evolved and mono-evolved *B. velezensis* populations based on a Bray-Curtis dissimilarity matrix. “Early” represents 1^st^ and 3^rd^ transfer, “Middle” represents 6^th^, 9^th^, and 12^th^ transfer, “Late” represents 15^th^ and 18^th^ transfer. **(E)** Bray-Curtis distance between the mutations of mono-evolved and co-evolved *B. velezensis* lineages over time. **(F)** Genealogy and genotype frequencies over time. The colors represent different genotypes, and the vertical area represents genotype frequency as determined by Lolipop. When a mutation arises in the background of another mutation, generating a new genotype, the new color arises in the middle of the existing genotype. Dominant genotypes are highlighted in certain colors, other genotypes are in gray. Mutations occurred in intergenic regions were excluded from this figure.

Nonmetric multidimensional scaling (NMDS) analysis based on Bray-Curtis dissimilarity revealed that in early evolution stages (1^st^ and 3^rd^ transfer), mono-evolutionary mutations were grouped together with co-evolutionary mutations (Fig 5D). However, the mutations grouped separately during the middle and late stages of evolution (6^th^ to 18^th^ transfer). Permutational multivariate ANOVA (PERMANOVA) showed that the mutation compositions between the two evolution groups differed significantly (*p* = 0.001, F = 9.555). Accordingly, the Bray-Curtis distance increased over time between mono-evolved and co-evolved *B. velezensis* populations (Fig 5E). The intra-group diversity did not significantly differ between mono-evolved and co-evolved *B. velezensis* lineages at each transfer stage. (Fig S5A & B). Using lolipop software package, we inferred the genotype and genealogy of each population. The linked genotypes and ancestry were identified based on allele frequency data and displayed as Muller plots (Fig 5F). This figure shows the breadth of genotype frequency with colors representing the presence of *spo0F*, *ywcC*, *comA*, *comP*, *degS*, and *baeL* mutations within the genotype. Regardless of the evolution treatment, *spo0F* mutations were detected in all lineages and subsequently rose in frequency, reflecting the parallel evolution to the TSB-rich environment. The biofilm-enhancing *ywcC* mutations were prominent across 5/8 mono-evolved lineages, which is consistent with the increased number of cells in the final mono-evolved biofilm populations (Fig 1C). Mutations causing rough morphotype (*comA*, *comP*, *degS*) were observed in all co-evolved lineages, but only in 5/8 of the mono-evolved lineages. Mutations in *comA* and *baeL* were exclusively found in co-evolved lineages and always occurred together within the same genotype.

## Discussion

In most environment, microorganisms do not evolve in isolation but instead, the species will evolve in a community context. Interspecies interaction have been shown to affect adaptive evolution to new environments ^9,12,18,36–38^. It remains unclear how dynamically changing interaction relationships are impacted by evolution. This study examined how *B. velezensis* adapt in isolation and in interaction with *P. stutzeri*, including differences in phenotypic, genotypic, and fitness characteristics.

Both mono-evolved and co-evolved lineages increased total cell density after 36 days of biofilm evolution (∼120 estimated generations). *B. velezensis* ancestor repeatedly diversified into slimy and rough morphotypes in the parallel evolved lineages. These morphotypes displayed phenotypic differentiations: planktonic growing capacity increased while growth rate and spore development decreased. Accordingly, several genes associated with sporulation or regulation of spore development have been identified in the genomes of evolved clones and metagenomes of evolved populations. Sporulation is an essential trait for surviving the harsh environment in the soil, which is the habitat of the focal species, however, they are dispensable in the liquid rich medium used for this study. In other domestication assays using rich medium, the loss of sporulation trait was also reported ^39,40^. *P. stutzeri* colony evolved from irregular to round due to a mutation in the *cpsA* gene encoding CPS biosynthesis protein. In *Pasteurella multocida* and *Vibrio vulnificus*, mutants lacking the CPS formed significantly more biofilms than wild type ^33,34^. The lack of CPS can influence the evolutionary strategy of *Klebsiella variicola* to spatially structured environments, impacting traits such as aggregation, virulence, and morphotype differentiation ^41–43^. Even though the evolved *P. stutzeri* isolate and the mutant did not enhance pellicle formation in isolation, they enhanced the formation of dual-species biofilms through overproduction of extracellular polysaccharides.

The interaction of the two ancestral species shifted from cooperation to ammensalism from 24h to 48h is remisicent of the dynamic interaction between yeast and a bacterium: at an early stage, yeast secreted bacterial growth promoter that block the transport of bacterial inhibitor and established commensalism; when the growth promoter became depleted and the growth inhibitor accumulated, their interaction shifted from commensalism to ammensalism ^14^. Based on these findings, it is important to consider the time effect when defining the type of speices interactions. The final interaction between *B. velezensis* and *P. stutzeri* is also a mix of mutualism and competition: the evolved *P. stutzeri* interacts mutually with rough isolates of *B. velezensis*, but it engages in ammensalism with the slimy isolates. Both biotic interaction and abiotic adaptation led to phenotypic diversification, although with distinct morphotype frequencies and fitness landscape under these two conditions. This observation corresponds to a previous biofilm evolution study by Røder and colleagues, although the wrinkled variants of *X. retoflexus* emerged in both in mono- and cocultures, while the selective pressure applied by *P. amylolyticus* increased the frequency of the variants ^10^,.

Among both mono- and co-evolved isolates, the slimy morphotype displayed increased pellicle formation, while the rough morphotype formed less floating biofilm. The colony morphotype change of slimy isolates could be reproduced by mutating the *spo0F* gene, while their increased pellicle formation could be attributed to frameshift mutations in *ywcC* gene. The *ywcC* locus has been identified as a hotpot gene for mutations when *B. subtilis* evolves towards enhance root colonization ^44,45^. Further mutations in *comA*, *comP*, *degS*, or *degQ* genes might counteract the effect of *ywcC* mutations and led to impaired formation of the rough isolates. Five co-evolved rough isolates, as well as four co-evolved lineages, were found to have mutations in the *baeL* gene, which involves in bacillaene synthesis. Co-evolving with *P. fluorescens* on tomato roots, *B. subtilis* displayed mutations in the *pksR* gene, which is also involved in synthesizing bacillaene ^44^. Bacillaene was reported to have bacteriostatic activity on *P. chlororaphis* ^46^. Mutation in *baeL* gene could be a strategy to minimize inhibitory effect on *P. stutzeri*. However, the lack of antagonistic activity against *P. stutzeri* by *B. velezensis* suggests that other mechanisms or pathways might be involved in this specific interaction. Similar to the smooth variants of *B. subtilis* observed during evolution for subsequent pellicle formation ^25^, the rough variants in this study could exploit the secreted biofilm matrix components of the other morphotypes to promote pellicle formation. Apart from that, the extracellular matrix produced the evolved *P. stutzeri* isolates could also promote the pellicle formation of rough variants. Even though pellicle biomass was highest in cocultures containing co-evolved slimy isolates, the ammensalism renders it evolutionary unstable and susceptible to replacement by weak but cooperative rough isolates. These results show that dual-species community can select for adaptive exploiters which are cooperators in the community. This finding is in line with the so-called Black Queen Hypothesis, the evolution of dependency through trait loss ^17^. The results in our study emphasize the need for considering interspecies interaction when exploring intraspecies adaptation.

In summary, this study reported that interspecies interaction alters the fitness landscape of evolved genotypes and selects for adaptive exploiters in partner populations. Currently, this controlled system serves as a model for investigating the coevolution of these two bacteria, laying the groundwork for future research into their coevolutionary dynamics in the rhizosphere environment and ultimately exploring their biological functions and impacts on plant health.

## Methods and materials

### Strains and media

*B. velezensis* strain SQR9 and *P. stutzeri* strain XL272 were the ancestral strains used in the evolution experiments ^20^. *B. velezensis* SQR9 was transformed with a plasmid containing a GFP marker, *P. stutzeri* XL272 was modified with a mini-Tn7 transposon containing a dsRed marker. Tryptone soya broth (TSB) (Sigma-Aldrich, CAT# 22098) was used for all experimental evolution and phenotypic characterization. Lysogeny broth (LB) was used during genetic engineering. Solid media contained 1.5% agar added to TSB or LB media. All strains were stored at −80 °C using overnight grown cultures in TSB or LB media supplemented with 25% glycerol before freezing the samples.

Start inoculum across the study was obtained by growing the cells overnight to exponential phase in TSB medium at 30 °C, 180 rpm shaken condition, the cells were spun down and diluted to OD_600_ of 1 in 0.9% NaCl buffer. For coculture, the start inoculum was prepared by mixing equal volumes of isolates.

### Experimental evolution

Approximate 10^6^ cells of *B. velezensis* SQR9 and *P. stutzeri* XL272 from independent overnight cultures were inoculated in non-adjacent wells of two independent 24-well cell culture plates (VWR, CAT# 10062-898) and incubated at 30 °C in TSB supplemented with 5 mg/l chloramphenicol without shaking. Monoculture of *B. velezensis* was used as control evolutionary treatment. Addition of chloramphenicol was to preserve the plasmid in *B. velezensis*. Since *P. stutzeri* cannot form pellicle, mono-evolution of *P. stutzeri* was not performed in this study. Every two days, mature pellicles from the first plate were harvested with sterile inoculation loop, disrupted and re-inoculated in two independent plates through a 1:100 dilution (20 μl in 2 ml of fresh TSB + chloramphenicol medium) according to the method described in literature ^2^. The remaining populations were mixed with 1 ml of 50% glycerol solution and preserved as frozen stocks (−80 °C). The second plate was imaged using an Axio Zoom V16 stereomicroscope (Carl Zeiss, Jena, Germany) equipped with a Zeiss CL 9000 LED light source and an AxioCam MRm monochrome camera (Carl Zeiss) equipped with HE eGFP (excitation at 470/40 nm and emission at 525/50 nm) and HE mRFP (excitation at 572/25 nm and emission at 629/62 nm) filter sets. After the 1st, 3rd, 6th, 9th, 12th, 15th, and 18^th^ transfer, 10% of the disrupted populations were sonicated (24*1s pulses at 20% amplitude with 1 s pause between the pulses), serial diluted and plated on TSB plates supplemented with either 5% NaCl for *B. velezensis* or 20 mg/l gentamicin for *P. stutzeri*. Colonies with divergent morphotypes were randomly selected and preserved at −80 °C after overnight cultivation in TSB medium and supplementation of the culture with 25% glycerol.

### Phenotypic characterizations

(1) Colony morphology assay. Colony morphologies were examined on TSB medium with 1.5% agar. The plates were dried under laminar airflow conditions for 40 min after solidifying. Glycerol stocks were streaked out and incubated at 30 °C until colonies were visible. Single colonies were re-streaked out on fresh plates and incubated at 30 °C. Photos were taken at 2-3 days for *B. velezensis* and 6 days for *P. stutzeri*.
(2) Pellicle morphology assay. 20 μl of the start inoculum was cultivated in 2 ml of TSB medium in a 24-well microplate (Fisher Scientific). The microplates were incubated statically at 30 °C for 48 h.
(3) Growth curve assay. 100 μl start cultures were inoculated to 1 ml TSB medium in a 48-well microplate and grown under shaken conditions at 30 °C in a Bioscreen C Pro Automated Microbiology Growth Curve Analysis System with OD_600_ measurements every 15 min. Each treatment contained five replicates.
(4) Spore observation and quantification. Pellicles formed after 2 and 7 days were collected and resuspended in 2 ml of PBS buffer with glass beads, then vortexed vigorously for 5 minutes to disrupt the pellicles into single cells and aggregates. These disrupted cells were observed using Phase Contrast and Darkfield Microscopy (Olympus CX41). Spores appear transparent, while vegetative cells and dead cells appear black. 1 ml of the disrupted suspension was heat-treated at 70 °C for 15 minutes to kill vegetative cells. Both the disrupted and heat-treated samples underwent sonication (30% amplitude, 30 cycles of 1 s sonication with 1 s rest) to further disaggregate cells. The total cell numbers and spore numbers were quantified using serial dilution plating and colony counting after incubation.
(5) Quantification of EPS in the pellicle matrix. Pellicles were grown in 24-well microplates as described before. The supernatant was removed by syringe, pellicles were suspended in 2 ml water. Cells were separated from pellicle matrix by vortexing vigorously and centrifugation (10,000 g, 2 min), and the supernatant was filtered through 0.22 μm pore size filter. Total polysaccharides in the filtrate were quantified by the phenol-sulfuric acid assay with glucose as standard ^47^.

### Pellicle complementation assay

The rough morphotypes (BP1R, BP6R) were supplemented with the pellicle matrix or supernatant of the other *B. velezensis* morphotype isolates. The matrix donor strains contained *B. velezensis* ancestor, two coevolved slimy isolates (BP1S, BP4S), and the evolved *P. stutzeri* (PE). 100 μl of the donor strains (OD_600_ ∼ 1) were inoculated in 6-well microplates containing 10 ml TSB medium, and cultivated at 30°C for 48 h. The underneath supernatant was removed by 10 ml sterile syringe, the pellicles were resuspended in 10 ml water and vortexed rigorously for 15 mins. For PE, the supernatant was collected instead. The collected pellicles were centrifuged at 10,000 g for 5 min, and filtered through 0.22 μm pore size filter. The pellicle matrix was mixed with equal volume of 2*TSB medium. Sterile water was mixed with 2*TSB as a control medium. 20 μl of the recipient strain was inoculated in these media in 24-well microplates and incubated statically at 30 °C for 48 h. Photos were taken for these pellicles. The supernatant was removed by sterile syringe, the pellicles were resuspended in 2 ml water and vortexed rigorously for 15 minutes. The pellicle suspension was transferred to 48-well plates and the biomass was determined by measuring the optical density at 600 nm. Each treatment contained 6 replicates.

### Construction of *P. stutzeri* gene knockout mutant

The oligonucleotide primers used in this study are listed in Table S1. Knockout mutant for the *cpsA* gene was generated by DNA recombination forced by I-SceI endonuclease followed by counter selection ^48,49^. Upstream and downstream fragments were amplified by appropriate primers with recognition sites of I-SceI, digested by restriction enzymes, assembled by ligase, and cloned into pSNW4 vector. The resulting plasmid was introduced into XL272 by triparental mating using pRK600, generating the integrant strain. The helper plasmid pQURE which encode the I-SceI endonuclease was introduced into the integrant strain by electroporation to force the double DNA recombination. Knockout mutant was validated by PCR and sequencing. The helper plasmid was cured by omitting the inducer compound from the medium.

### Cell numbers quantification in pellicles

Cell numbers of both species in pellicles were quantified by absolute qPCR using the same primers described in ^20^. For tracking the cell number changes throughout evolution, 500 μl of population from 1^st^, 3^rd^, 6^th^, 9^th^, 12^th^, 15^th^, and 18^th^ transfer was taken out from the frozen stocks. Each population only contained one sample at each transfer. To compare the pellicle biomass of evolved isolates, samples were collected from pellicles grown in TSB medium for 48 h in 24-well plates. Each treatment included three biological replicates. Genomic DNA was extracted with EURx Bacterial & Yeast Genomic DNA Kit (CAT# E3580). qPCR was performed with Agilent’s Real Time PCR instrument Mx3005P. Reaction components are as follow: 7.2 μl H_2_O, 10 μl 2× Luna Universal qPCR Master Mix (NEB, CAT# M3003), 0.4 μl 10 μM of each primer and 2 μl template DNA. The PCR programs were carried out under the following condition: 95 °C for 1 min, 40 cycles of 95 °C for 5 s, 60 °C for 45 s, followed by a standard dissociation curve segment. Each sample was run in triplicates.

### Evolved clone genome re-sequencing and analysis

Sixty-nine *B. velezensis* evolved clones and sixteen *P. stutzeri* evolved clones were selected for genome resequencing. Four ancestors of each species were also re-sequenced to exclude common mutations arose from historical lab domestication instead of this study. The strain list is included in Table S2. Genomic DNA of selected strains were extracted using the EURx Bacterial & Yeast Genomic DNA Kit from overnight cultures. Paired-end fragment reads (2*150 bases) were generated using an Illumina HiSeq/Novaseq instrument. Raw reads were analyzed by Cutadapt (V1.9.1) to remove adaptors, primer sequences, content of N bases more than 10%, and bases of quality lower than 20. Clean reads were mapped to the reference genomes (CP046538.1 and CP009679.1) using BWA (V0.7.17). Mapping results were processed by Picard (V1.119) to remove duplication. Variants were called using GATK (V3.8.1) software and annotated using Annovar (V21 Apr 2018). Data on mutation frequencies for each strain sequenced are provided in Table S2. Mutations in were then filtered in R to remove mutations (1) with low frequency (<95%), (2) synonymous SNP, (3) intergenic mutations. Subsequently, mutations identified in the ancestral genomes were filtered. The resulting table was visualized using the R *pheatmap* package.

### Whole-genome population sequencing and analysis

Whole populations were sequenced at 1^st^, 3^rd^, 6^th^, 9^th^, 12^th^, 15^th^, and 18^th^ transfer. All sixteen populations (eight from mono-evolution, and eight from co-evolution) and ancestor strains were sequenced. Genomic DNA were extracted from 1 ml of frozen populations using the EURx Bacterial & Yeast Genomic DNA Kit. The sequencing libraries were prepared using MGIEasy Fast FS DNA Library Prep Set (MGI Tech). Paired-end fragment reads (150bp ×2) were generated on a DNBSEQ-Tx sequencer (MGI Tech) following the manufacturer’s procedures. All population samples were sequenced with ultrahigh depth data to ensure > 300 × depth coverage for each species in evolved and co-evolved conditions. Raw sequencing data have been deposited into CNGB Sequence Archive (CNSA) of China National GeneBank DataBase (CNGBdb) with accession number CNP0003899.

Raw data were filtered using SOAPnuke^50^ (version 1.5.6) to remove low quality reads: reads including more than 50% of bases with quality lower than 12, reads including more than 10% of unknown base “N”, and reads containing adaptor contamination. For the co-evolved populations, the clean data were mapped to the *Bacillus velezensis* SQR9 genome (GenBank accession number CP006890.1) and *Pseudomonas stutzeri strain* XL272 genome (GenBank accession number CP046538.1) using bowtie2 with the parameters used in breseq^51,52^ (version 0.35.7), and the clean data of the mono-evolved populations were only mapped to the SQR9 genome. Then reads which mapped to SQR9 genome or XL272 genome were extracted from bam file using samtools^53^ (version 1.12), respectively. To ensure the similar variants calling sensitivity, we normalized the mapped data to 300 bp depth using seqtk (version 1.3-r107-dirty) (https://github.com/lh3/seqtk) for each species in each population sample for mutation calling. Mutations were called using breseq (version 0.35.7) with the default parameters and a -p option for population samples. The default parameters called mutations only if they appeared at least 2 times from each strand and reached a frequency of at least 5% in the population. The resulting mutations were provided in Table S3. We then filtered the mutations that found in the ancestor strains and those that never reached a cumulative frequency of 5% within a lineage. The resulting mutation list was subjected to analysis and visualization. The Jaccard similarity coefficient was calculated using the following formula: the number of mutated genes shared by both lineages divided by the total number of mutated genes in both lineages. NMDS based on a Bray-Curtis dissimilarity matrix was performed and plotted using the R vegan package to explore the differences in the composition of mutations. PERMANOVA was conducted to evaluate the effects of evolution treatment on the composition of mutations by using the R vegan package. We filtered out mutations that occurred only once in each lineage and those in intergenic regions, then visualized the mutational dynamics using Muller plots. Muller plots were generated using the Lolipop package^54^ (version 0.6) (https://github.com/cdeitrick/lolipop) using default parameters. This package predicts genotypes and populations based on shared trajectories of mutations over time and test their probability of nonrandom genetic linkage. Successive evolution of genotypes, or nested linkage, is identified by a hierarchical clustering method. Then Muller plots were manually colored by genotypes.

## Data availability

Data used in the main figures were provided in Table S4. Data used in the supplementary figures were provided in Table S5.

## Acknowledgements

This work was financially supported by the National Nature Science Foundation of China (42307173), the China Postdoctoral Science Foundation (2023M747140). XS was supported by China Scholarship Council fellowship (201906850104), Postdoctoral Fellowship Program of CPSF (GZB20230309), Jiangsu Funding Program for Excellent Postdoctoral Talent (2023ZB250). ÁTK was supported by the Novo Nordisk Foundation within the INTERACT project of the Collaborative Crop Resiliency Program (NNF19SA0059360), the Danish National Research Foundation (DNRF137) for the Center for Microbial Secondary Metabolites, and a start-up fund from Institute of Biology Leiden. GH was supported by the China National GeneBank (CNGB). JX was supported by a China Scholarship Council fellowship.

## Author contributions

XS, ÁTK, ZX, QS designed the study, XS, JX, YL, LT performed the experiments. XS, GH, YM analyzed the data and created the figures. XS wrote the first draft of the manuscript, ÁTK, ZX, NZ, WX, RZ, QS revised the manuscript.

