## Supplementary material for "Interspecies interaction reshapes the fitness landscape of evolved genotypes": Fig S1 to S5

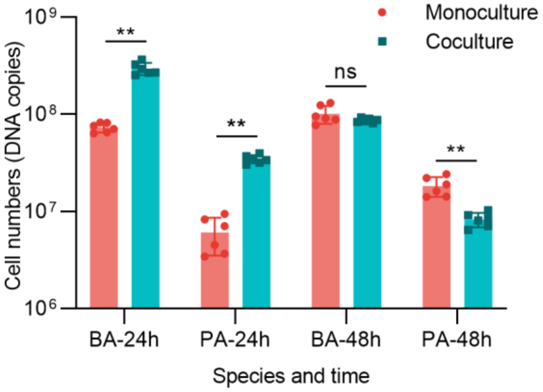


**Fig. S1 Cell numbers quantification in pellicle.** Dual-species pellicle containing ancestral isolates of 24h and 48h. “BA” represents *B. velezensis* ancestor, “PA” represents *P. stutzeri* ancestor. “**” indicates significant differences ($p$<0.01) based on t test.

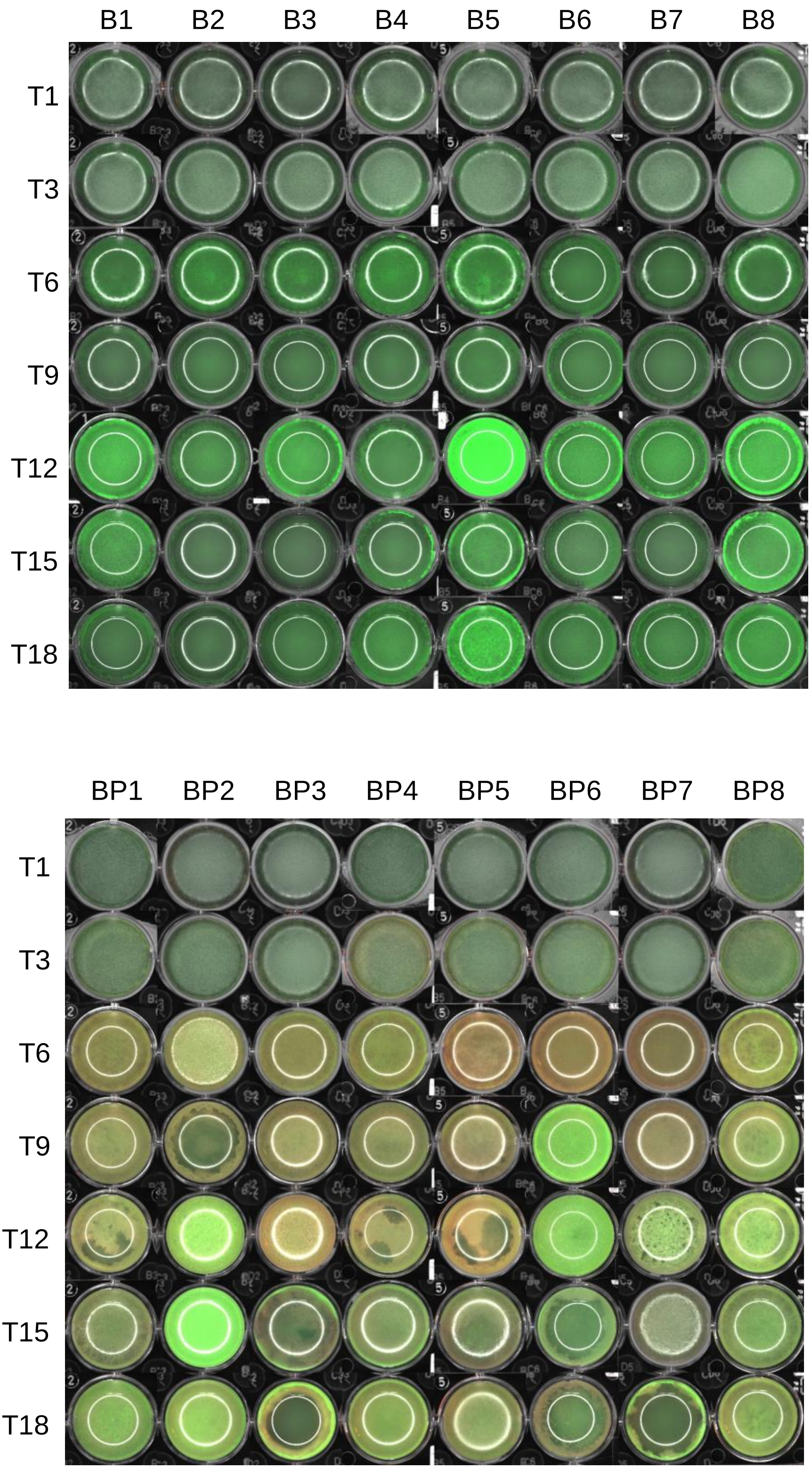


**Fig. S2 Phenotype of evolving pellicle populations. (A)** Mono-evolution. **(B)** Co-evolution. *B. velezensis* were labeled green, *P. stutzeri* were labelled red, the overlay were yellow or orange depending on the proportion of the two species.

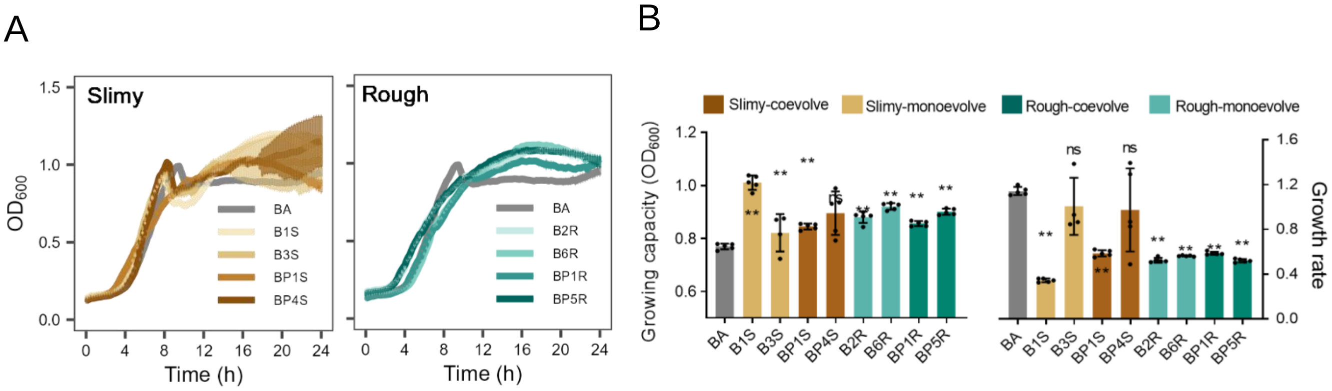


**Fig S3. Phenotypic characterization of *B. velezensis* evolved variants. (A)** Growth curve. n = 5. **(B)** Growing capacity and growth rate. Growing capacity represents the maximum population size. “ns” indicates no significant difference with ancestor, * $p$ <0.05, ** $p$ <0.01 by t test.

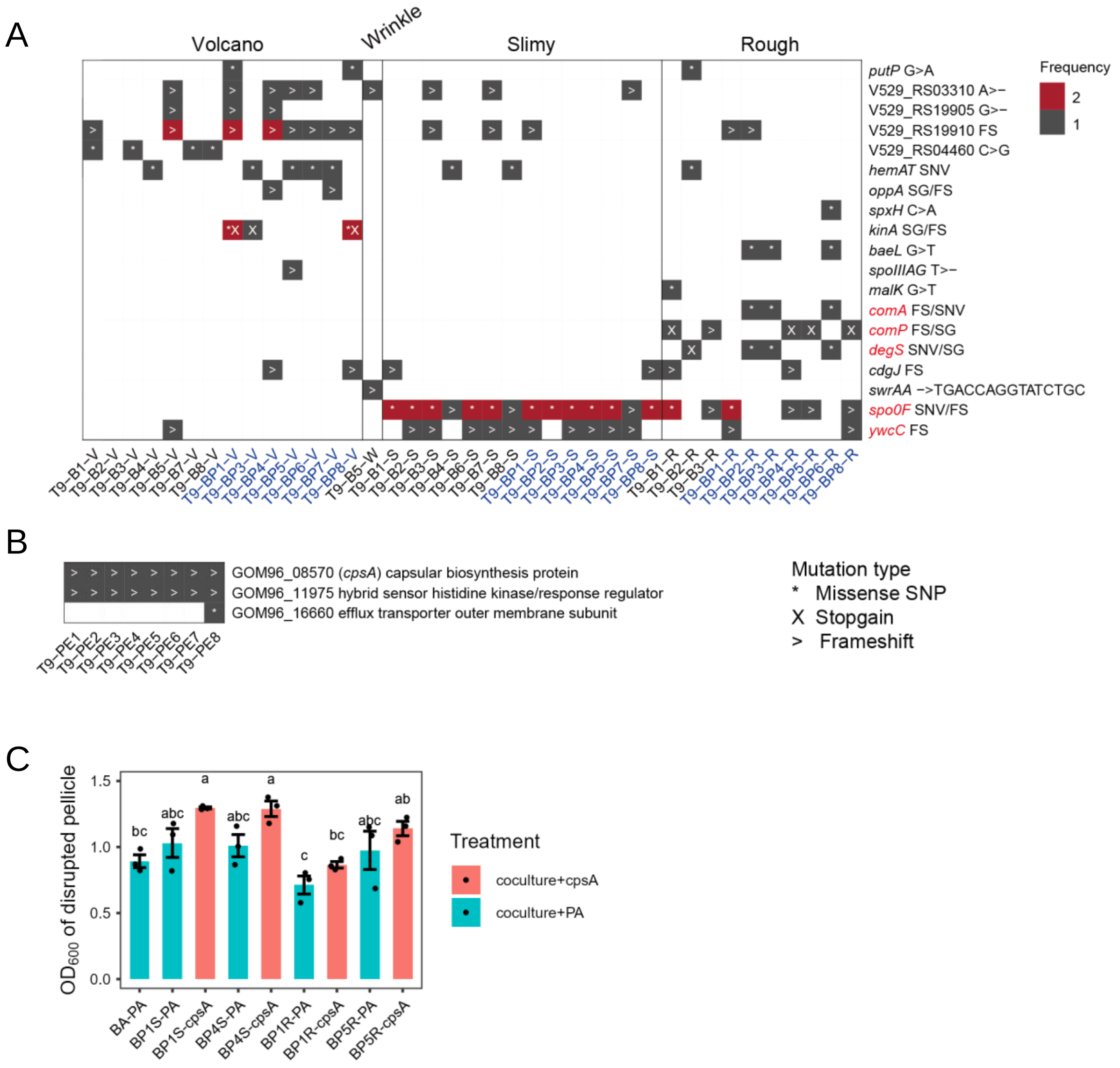


**Fig. S4 Mutations identified in evolved clones from 9th transfer. (A)** *B. velezensis* SQR9 mutations**.** V abbreviates for volcano, W abbreviates for wrinkle, S abbreviates for slimy, R abbreviates for rough **(B)** *P. stutzeri* XL272 mutations. PE abbreviates for evolved *P. stutzeri*. For both (A) & (B), "n" indicated the population. "*" and SNP refer to missence single nucleotide polymorphism, ">" and FS refers to frameshift mutation, "X" and SG refers to stopgain mutation. All the isolates shown above were isolated from the 9th transfer. (C) Pellicle quantification by

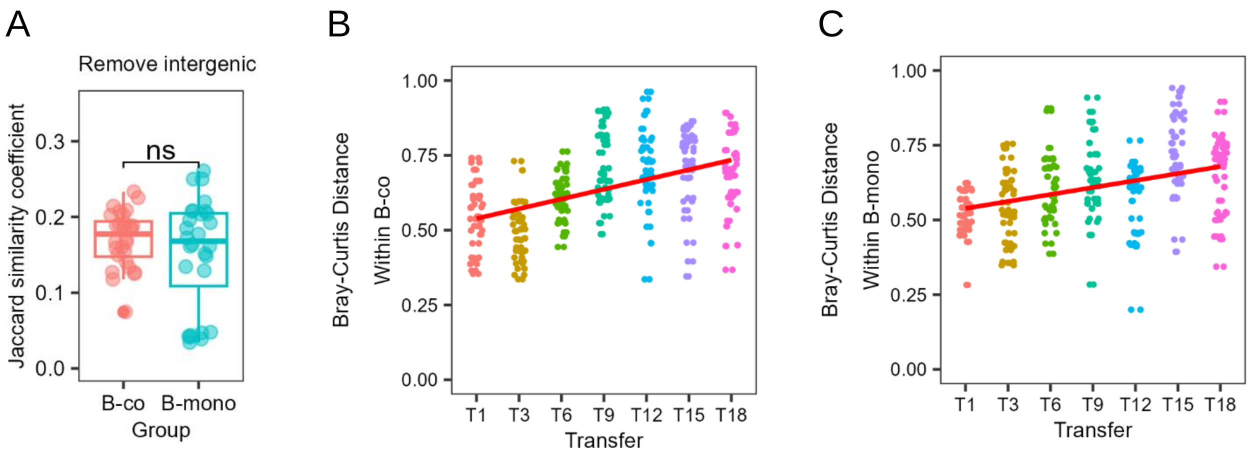


**Fig. S5 Analysis of population sequencing. (A)** Degres of parallelism within each estimated by Jaccard indexgroup after removing the mutations occurred in intergenic regions. **(B-C)** Bray-Curtis distance within the mutations of co-evolved *B. velezensis* lineages (B) and mono-evolved lineages (C) over time.
